# Multiomic analysis of non-glaucomatous human trabecular meshwork cells

**DOI:** 10.64898/2026.07.28.740161

**Authors:** Anh H. Pham, Isabella G. Moceri, Zachary Batz, Nivedita Singh, Avinash Soundararajan, Katelyn Kane, Tutut Nurjanah, Emine Bilir, Rong Du, Yiqin Du, Milton A. English, Jennifer A. Faralli, Samuel Herberg, Stephanie How, Ruminder Preet Kaur, Susanna Li, Suhani Patel, Padmanabhan P. Pattabiraman, Carl M. Sheridan, Anand Swaroop, Ying Ying Sun, Neeru A. Vallabh, Sanjoy K. Bhattacharya, Donna M. Peters, Kate E. Keller

## Abstract

Glaucoma is an irreversible blinding disease that affects millions of individuals worldwide. Elevated intraocular pressure (IOP), regulated by the trabecular meshwork (TM) in the anterior eye, is the only modifiable risk factor. Human TM cells can be cultured from donor eyes, providing a precious resource for studying factors that induce or prevent glaucoma. The goal of this study was to produce datasets that define the molecular profile of human non-glaucomatous TM cells. Using 18 human TM cell strains cultured from non-glaucomatous individuals deposited from seven laboratories in the USA and UK, this study used transcriptomic, proteomic, lipidomic, and metabolomic analyses to characterize the molecular content of TM cells. The data herein provides the most comprehensive multiomic analyses of human TM cells to date and will be a useful resource for researchers and clinicians in the TM and glaucoma fields.

## Background and Summary

The trabecular meshwork (TM) is an important tissue in the conventional aqueous humor outflow pathway of the eye. The TM is located in the anterior segment, in the angle between the cornea and iris. The tissue functions as a biological filter and is crucial for determining intraocular pressure (IOP) by regulating outflow of aqueous humor from the anterior chamber into Schlemm’s canal (SC), which then drains out via the episcleral venous plexus and into the bloodstream.^1^ Structural and functional dysfunction of TM leads to increased IOP, an endophenotype associated with primary open-angle glaucoma (POAG). Increased IOP exerts mechanical stress at the back of the eye, leading to retinal ganglion cell death, optic nerve degeneration, and irreversible blindness.

The TM is composed of three layers. The first two layers comprise a mesh-like network of trabecular beams, composed of extracellular matrix (ECM) and surrounded by a monolayer of endothelial-like cells.^2^ The third layer, known as the juxtacanalicular region, is composed of cells embedded in an abundant ECM and connected to one another via cellular processes. These TM cells have characteristics of both fibroblasts and smooth muscle cells and contribute to the tissue’s contractile properties that move fluid through the TM. Cells in the TM also exhibit phagocytic properties and play an important role in engulfing cellular debris from aqueous fluid before it enters and clogs the outflow tract. TM cells detect and respond to changes in IOP to initiate a homeostatic response that maintains IOP within a narrow range. This is an active area of research that involves a complex process of mechanotransduction signaling events^3^ and the regulation of ECM assembly and turnover.^4^ Thus, a better understanding of the biological properties of healthy TM cells is important for maintaining normotensive IOP and normal vision.

Comparisons between glaucomatous and non-glaucomatous TM tissues have identified genes and proteins implicated in POAG pathogenesis. Key genes/pathways harboring detrimental changes include elevated TGFβ signaling,^5,6^ concomitant synthesis of excess ECM,^4,7,8^ abnormal Wnt signaling,^9^ deleterious changes to the actin cytoskeleton that results in thicker, abundant actin stress fibers, alterations to integrin levels,^10^ and activation of mechanosensitive YAP/TAZ signaling.^11,12^ These changes contribute to the increased biomechanical stiffness observed in glaucomatous TM.^3^ Most of the studies have used the standard consensus recommendations of at least 3 biological replicates of TM cells or tissue, with several using additional biological replicates. Moreover, no study has compared biological differences among non-glaucomatous TM cells. This is important because it will reveal new information regarding the recommended number of non-diseased cell strains that should be included in a study to partially compensate for human genetic variability. Importantly, multiomics has not been performed on non-glaucomatous TM cells.

In this study, seven laboratories from across the US and the UK collaborated to provide 18 non-glaucomatous TM cell strains for analysis by transcriptomics, proteomics, lipidomics, and metabolomics. This is a first-of-its-kind study that provides the research community with rich datasets of non-glaucomatous TM cells. Notably, transcriptomic and proteomic data remained largely consistent across depositing laboratories, supporting that these datasets represent comprehensive molecular profiles of a TM cell. The lipid and metabolomics datasets show that age has a major influence on the lipid and metabolite profiles of TM cells. Furthermore, antidepressants and plastics were identified as metabolites found in certain cell strains, indicating that systemic medications and environmental exposure metabolites are retained in TM cells over several passages in cell culture media. The databases reported herein of genes, proteins, lipids, and metabolites can be a useful resource for researchers in the TM and glaucoma fields.

## Methods

### Ethics Statement

The Institutional Review Boards from each of the laboratories reviewed the protocols used to culture TM cells from human cadaver eyes and provided written confirmation that it is “not research involving human subjects” (Oregon Health & Science University, STUDY00025374; SUNY Upstate Medical University, IRB#1211036; University of Wisconsin minimal risk research, #2009-1274; University of Miami, IRB#20240793; University of South Florida; Indiana University School of Medicine, 1911117637). There were no living human participants in the study. All studies adhered to the Declaration of Helsinki regarding the use of human tissue for medical research.

### Trabecular meshwork cell culture

Eighteen human primary TM cell strains were obtained from 7 laboratories across the USA and from the UK. TM cells were isolated from human cadaver eyes procured from local eye banks: VisionGift (Portland, OR); Liverpool Research Eye Bank (Liverpool, UK); Lions World Vision Institute (Tampa, FL); and Beauty of Sight Eye Bank (Miami, FL), or from deidentified corneal rims obtained after surgical keratoplasty procedures. Demographics, tissue source, cause of death and clinical background of the human donors are listed in **Table 1**. Most of the cell strains were from Caucasian donors. However, information regarding race is sometimes incorrect or is unknown. The medical background is also extremely limited or unknown. The 18 TM cell strains used for the omics analyses were cultured from donor eyes from 7 females and 11 males. Four of the donor tissues were under 40 years old, seven were between the ages of 40 and 60 years, and seven were over 60 years old.

**Table 1.** Donor information and demographics for the 18 TM cell strains.

| Sample ID | PI | Original Donor ID | Age (Yr) | Sex | Tissue source | Cause of Death (if known) | Relevant clinical background (if known) |
| --- | --- | --- | --- | --- | --- | --- | --- |
| MC1 | KEK | 2018-1233 | 53 | Male | whole globe | Lung cancer w/metastases | Steroids, radiation therapy, antibiotic therapy |
| MC2 | KEK | 2020-0984 | 69 | Male | whole globe | Advanced Parkinsons and dementia | Hypertension, congestive heart failure |
| MC3 | KEK | 2022-1149 | 71 | Female | whole globe | Respiratory failure, severe multivessel coronary disease | Dementia, End stage renal disease, non-ST elevation myocardial infarction, aphasia, cardiomyopathy, severe mitral valve stenosis |
| MC4 | KEK | 2021-1493 | 74 | Female | whole globe | Subdural bleed, chronic myelomonocytic leukemia | Breast cancer, rheumatoid arthritis, osteoarthritis, endometriosis |
| MC5 | SHe | HTM 14 | 50 | Female | corneal rim | Unknown | Donor corneal rim obtained after surgery (PKP); medical history unknown |
| MC6 | SHe | HTM 19 | 34 | Male | corneal rim | Unknown | Donor corneal rim obtained after surgery (PKP); medical history unknown |
| MC7 | SHe | HTM 36 | 56 | Female | corneal rim | Unknown | Donor corneal rim obtained after surgery (DSAEK); medical history unknown |
| MC8 | YD | TM 165 | 57 | Female | corneal rim | Intracerebral hemorrhage (ICH) | Toe surgery, hypertension (no medication) |
| MC9 | YD | TM 214 | 61 | Female | corneal rim | COPD | No ocular history |
| MC10 | YD | TM 181 | 64 | Male | corneal rim | Anoxic brain injury / inhalation injury | Hypertension, coronary heart disease, myocardial infarction, coronary artery bypass grafting (unknown medications) |
| MC11 | DMP | #2354 | 25 | Male | corneal rim | Unknown | Donor corneal rim obtained after surgery; medical history unknown |
| MC12 | DMP | #2355 | 35 | Male | corneal rim | Unknown | Donor corneal rim obtained after surgery; medical history unknown |
| MC13 | PPP | HTM L-3 | 49 | Female | corneal rim | Unknown | Medical history unknown |
| MC14 | PPP | HTM L-4 | 68 | Male | corneal rim | Unknown | Medical history unknown |
| MC15 | AHP | 231011A | 83 | Male | Corneal rim | Pancreatic cancer, shortness of breath | Donor corneal rim obtained after surgery (PKP). No ocular history |
| MC16 | AHP | 2396E | 42 | Male | Corneal rim | Overdose | Donor corneal rim obtained after surgery (PKP). No ocular history |
| MC17 | CMS | NH 5034 | 29 | Male | Whole globe | Unknown | No Ocular history |
| MC18 | AHP | 2396D | 51 | Male | Corneal rim | Atherosclerotic cardiovascular disease | Donor corneal rim obtained after surgery (PKP). No ocular history |

KEK, Kate E. Keller; SHe, Samuel Herberg; YD, Yiqin Du; DMP, Donna M. Peters; PPP, Padmanabhan P. Pattabiraman; AHP, Anh H. Pham; CMS, Carl M. Sheridan. PKP = penetrating keratoplasty; DSAEK = Descemet Stripping Automated Endothelial Keratoplasty.

Tissue dissection procedures varied by donor source: either whole globe or corneal rim (**Table 1**). Some laboratories (KEK, SHe) dissected the TM tissue, digested with a collagenase, washed, and placed into fresh media for 2-3 weeks until the cells were confluent. Other laboratories (CMS) cut the TM into approximately 3-5 mm length sections and placed in a 6-well plate or 25 cm^2^ flask. Culture medium was added gently to cover the tissue, which was left undisturbed for 7 days to allow attachment and outgrowth of TM cells. To grow TM cells from corneal rims (PPP, AHP, DMP, YD), TM tissue was stripped from the corneal rim and placed in a gelatin-coated plate, sandwiched between a coverslip.

Culturing conditions varied by group, which is summarized in **Table 2**. For this study, TM cells were used between passages 3 and 5.

**Table 2.**
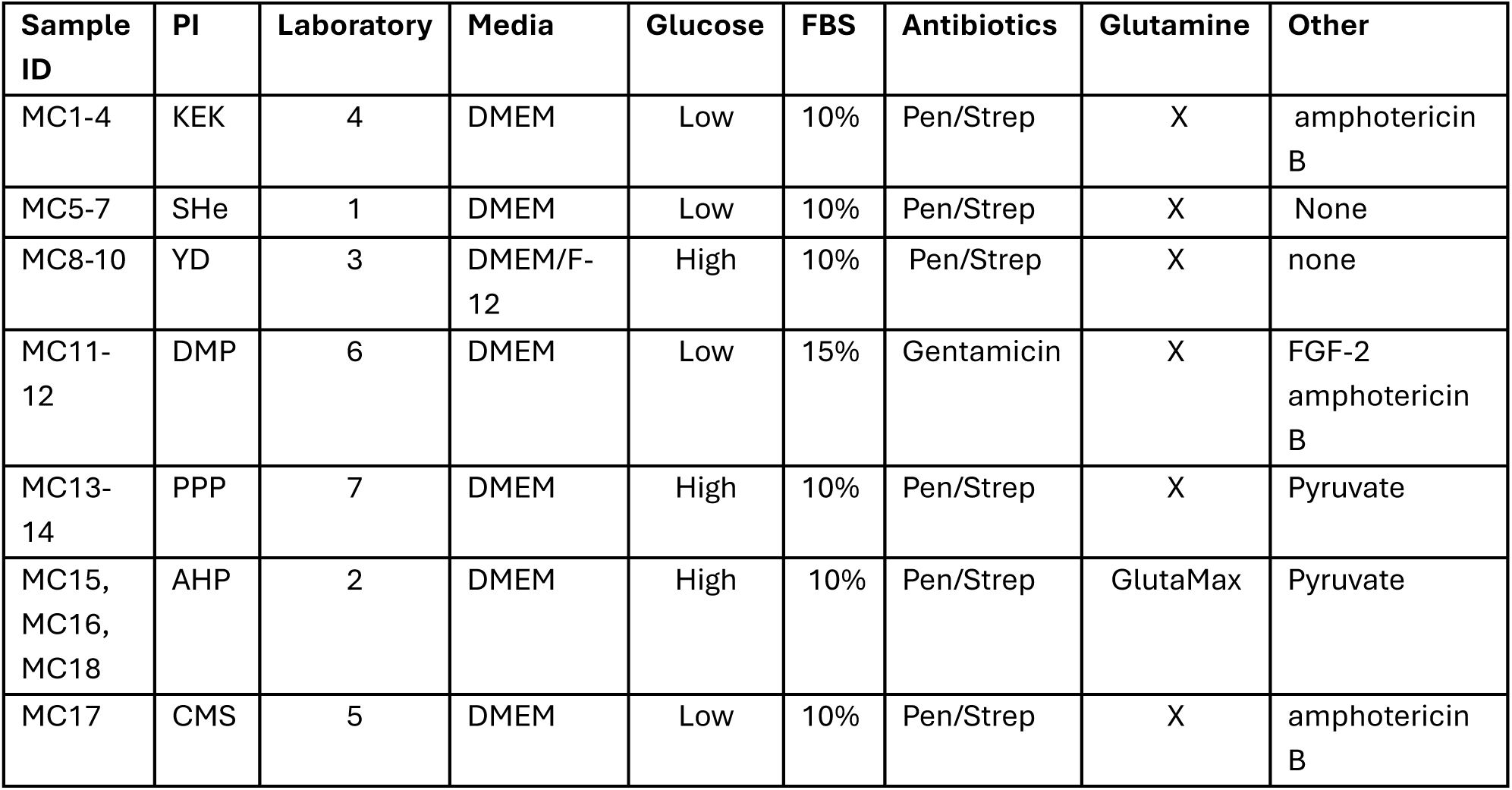
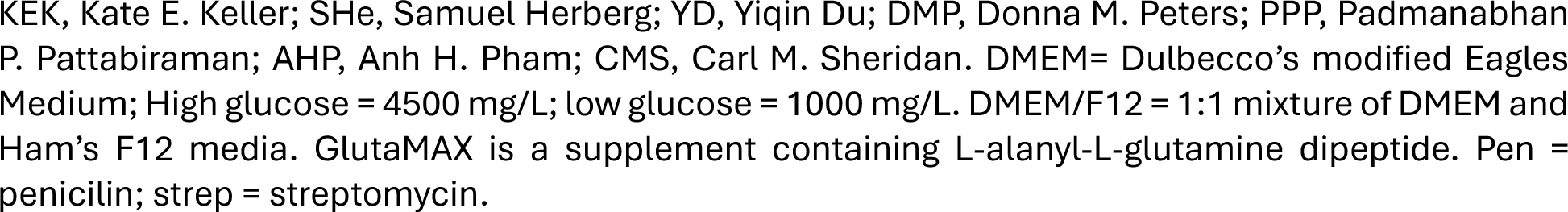
Cell culture conditions for each of the 18 TM cell strains.

| Sample ID | PI | Laboratory | Media | Glucose | FBS | Antibiotics | Glutamine | Other |
| --- | --- | --- | --- | --- | --- | --- | --- | --- |
| MC1-4 | KEK | 4 | DMEM | Low | 10% | Pen/Strep | X | amphotericin B |
| MC5-7 | SHe | 1 | DMEM | Low | 10% | Pen/Strep | X | None |
| MC8-10 | YD | 3 | DMEM/F-12 | High | 10% | Pen/Strep | X | none |
| MC11-12 | DMP | 6 | DMEM | Low | 15% | Gentamicin | X | FGF-2<br>amphotericin B |
| MC13-14 | PPP | 7 | DMEM | High | 10% | Pen/Strep | X | Pyruvate |
| MC15, MC16, MC18 | AHP | 2 | DMEM | High | 10% | Pen/Strep | GlutaMax | Pyruvate |
| MC17 | CMS | 5 | DMEM | Low | 10% | Pen/Strep | X | amphotericin B |

KEK, Kate E. Keller; SHe, Samuel Herberg; YD, Yiqin Du; DMP, Donna M. Peters; PPP, Padmanabhan P. Pattabiraman; AHP, Anh H. Pham; CMS, Carl M. Sheridan. DMEM= Dulbecco's modified Eagles Medium; High glucose = 4500 mg/L; low glucose = 1000 mg/L. DMEM/F12 = 1:1 mixture of DMEM and Ham's F12 media. GlutaMAX is a supplement containing L-alanyl-L-glutamine dipeptide. Pen = penicilin; strep = streptomycin.

Four tubes were collected per cell strain. Cells were harvested by washing twice in 1x phosphate buffered saline (PBS) and then dissociated with 0.05% trypsin-EDTA. After transferring to a tube, the cells were pelleted by centrifugation (1500g for 5 minutes) and the supernatant was removed. The cell pellet in the tube was frozen in an isopropanol bath and then stored at -80 °C. Each contained approximately 300-400,000 cells per tube. All tubes were subsequently shipped to Dr. Bhattacharya’s laboratory for proteomic, lipidomic, and metabolomic analyses, while one tube of each cell strain was subsequently shipped to Dr. Swaroop’s laboratory for transcriptomic analyses. An overview of the cell culture and ‘omics workflow is shown in **Figure 1**.

**Figure 1.**
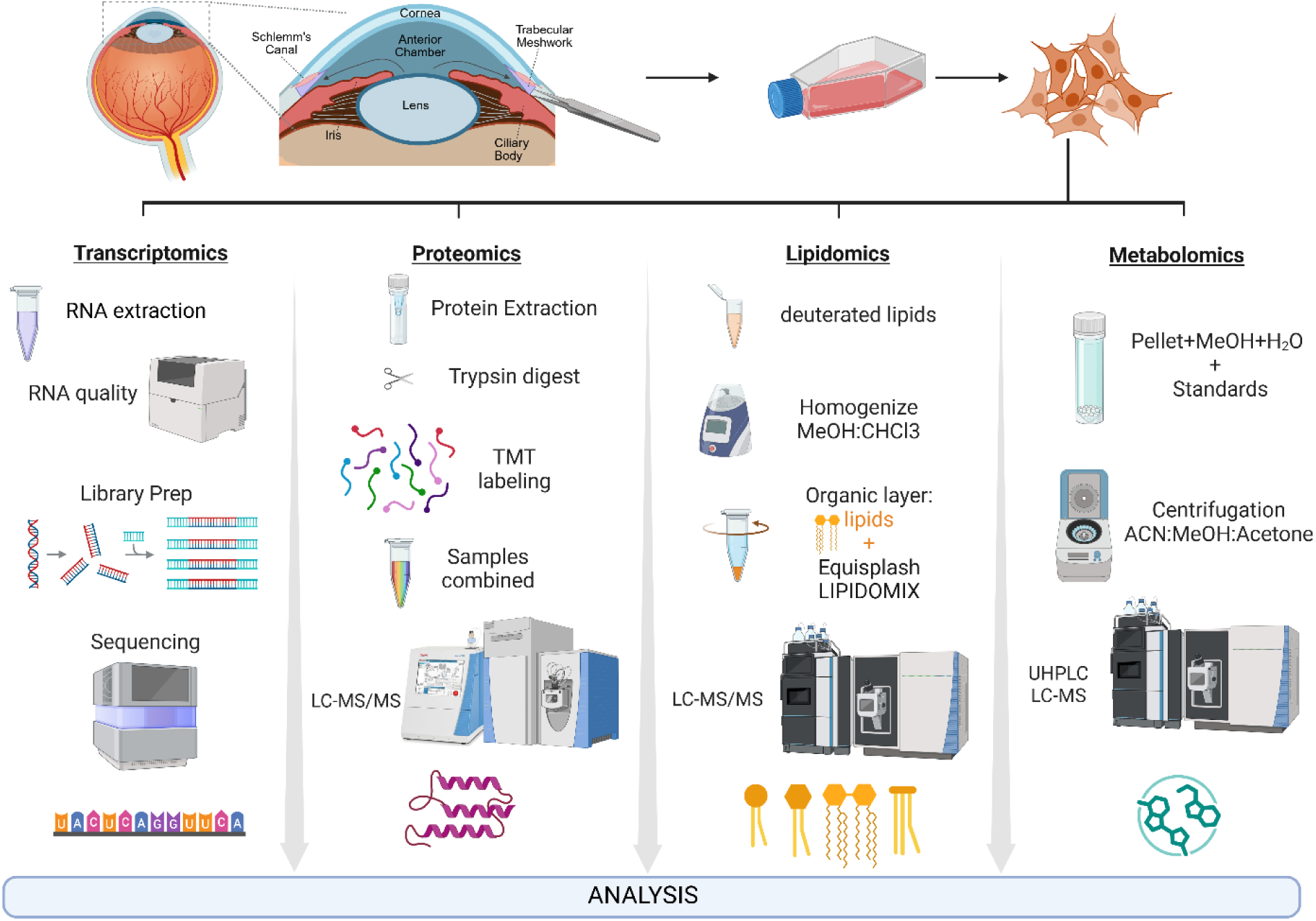
Trabecular meshwork cell culture and ‘omics workflow. The location of the TM is within the anterior portion of the eye. Human TM tissue was dissected and placed in cell culture. After primary cells reached confluence, they were harvested into four vials by the individual laboratories supplying the cells for transcriptomic, proteomic, lipidomic, and metabolomic analyses. The schematic shows the workflow for each of the omics.

### Total RNA-seq methods

Total RNA-seq was performed on 17 primary TM cell strains as previously described.^13^ Sample MC4 was not analyzed because it was lost in shipment. Total RNA was isolated using Qiagen RNeasy Mini Kit (Germantown, MD, USA; cat# 74104) and the RNA quality was assessed using Agilent TapeStation (Santa Clara, CA, USA). Given the partial degradation of the samples (RIN < 7), rRNA-depleted total RNA sequencing was selected over poly(A) selection to ensure uniform transcript coverage and minimize 3’ coverage bias.^14,15^ Libraries were prepared from 10ng of total RNA using the Illumina Stranded Total RNA Prep Ligation with Ribo-Zero Plus kit (San Diego, CA, USA; cat# 20040525). Libraries were sequenced on an Illumina NextSeq 2000 and analyzed as described previously.^16^ In brief, reads were trimmed using Trimmomatic v0.36 and pseudoaligned to the human reference genome GRCH38.p7 with the Ensembl v111 annotation using kalliso v0.45.0. Transcript-level counts were summarized to the gene level using tximport v1.30.0 then converted to counts per million, TMM-normalized, and tested for differential expression using the log likelihood ratio test in edgeR v4.0.3. PCA plots were generated using PCAtools v2.14.0. Volcano plots were generated with EnhancedVolcano.

### Protein extraction methods

The TM samples were processed following established protocols.^13^ Protein extraction was carried out using the EasyPep^TM^ Magnetic MS Sample Prep Kit, 96 reactions (catalog no. A57867, Thermo Fisher Scientific, Waltham, MA) following manufacturer’s recommendations. Prior to extraction cells were subjected to three cycles of freeze thaw (-80°C and 25°C each for 10 minutes duration) and sonication using a sonicator (ELMA ULTRASONICS Ultrasonic Cleaner, Model P30H; Elma Schmidbauer GmbH, Singen, Germany) for 5 minutes at 25°C. Prior to homogenization, three Regen III synthetic peptide standards— LLO (GYKDGNEYI), SEB (KKKVTAQELD), and CFP (EISTNIRQAGVQYSR) were spiked into the extraction buffer at 36 μM each to serve as internal controls for extraction efficiency. Following homogenization, lysates were transferred to microcentrifuge tubes and centrifuged at 12,000 × g for 10 minutes at 4°C. The resulting supernatant was carefully collected into fresh tubes for downstream processing. Protein concentration was estimated by dot blot densitometry and measured according to a BSA standard curve on ImageJ software. All samples were normalized to 50 μg/μL. Each of the 18 TM samples was labeled with a unique channel from the 18plex TMTpro isobaric labeling kit (catalog no. A52045, Tandem Mass Tag; ThermoFisher Scientific). After quenching the reaction, labeled samples were then pooled and dried via CentriVap Vacuum Concentrator (catalog no. 7810014, LABCONCO, Kansas City, MO). To the pooled sample, 2 μL of two Regen II human peptide standards A1315 (DRV[U-¹³C₅,¹⁵N]YI[U- ¹³C₆,¹⁵N]HPFHL) and HH4B (SGRGKGGKGLGKGGAKRHRKVLRGGK-Biotin) were added at 54 μM each as post-extraction ionization controls, together with 1 μL BSA and 47 μl 2% ACN. Both Regen III and Regen II peptide standards (>98% purity) were custom synthesized by AnaSpec (Fremont, CA, USA).

### Lipid extraction methods

Lipid extractions were performed using the modified Bligh and Dyer method^17^ as described previously.^18,19^ Briefly, TM cells were extracted in chilled 1:1 chloroform-methanol mixture. Uniformity of extraction was ensured by adding an internal standard composed of multiple deuterated lipid classes, EquiSPLASH^TM^ LIPIDOMIX® (Avanti Polar Lipids, Alabaster, AL, cat#330731) before cell homogenization. Samples were homogenized using the Precellys Homogenizer (Bertin Corp, Rockville, MD) system. Phase separation was obtained with the addition of chloroform:methanol:water (40:40:20 v/v) to the samples followed by high-speed spin (17,800 g) for 30 min at 4 °C. The organic phase containing lipids was transferred to a new tube and concentrated in a CentriVap Vacuum Concentrator (LABCONCO). Samples were regularly flushed with argon gas to prevent oxidation. The aqueous phase containing proteins was stored at −80 °C for protein quantification by the Bradford’s method and lipid/protein ratio. All extractions and subsequent handling were carried out using glass vials to minimize plastic contaminants in the samples.

### Metabolomic extraction methods

Metabolomic profiling was performed as previously described.^20,21^ The cell pellets were stored at -80 °C until ready for extraction. Metabolite extraction from pellets was carried out quickly while keeping the cells on dry ice to prevent degradation. Pellets were transferred to 0.5 mL Soft Tissue Lysing Kit Precellys tubes containing beads. 84 μL of chilled 1:1 methanol:H_2_O was added with four internal pre-extraction standards (all from Cayman chemicals, Ann Harbor, MI): 5 μl of 1 mg/ml Caffeine- ^13^C_3_ (catalog no. 27972), 5 μl of 1 mg/ml D-(+)-Glucose-^13^C_6_ (catalog no. 26707), 5 μl of 1 mg/ml Oleic Acid-d17 (catalog no. 9000432), and 1 μl of 5 mg/mL L-Isoleucine-^13^C_6_ (catalog no. 9004090). Tissues were homogenized using Precellys 24 Touch and homogenate was transferred to a microcentrifuge tube and centrifuged at 18000xrcf for 20 min at 4 °C. Supernatant was collected and the remaining pellet was transferred to Precellys Lysing Kit tube. 84 μL of chilled 8:1:1 Acetonitrile:Methanol:Acetone was added to the residual pellet in addition to the rest of the pre-extraction internal standards (all from Cayman chemicals): 5 μl of 1 mg/ml Caffeine-^13^C_3_ (CRM), 5 μl of 1 mg/ml D-(+)-Glucose-^13^C_6_, 5 μl of 1 mg/ml Oleic Acid-d17, 1 μl of 5 mg/mL L-Isoleucine-^13^C_6_. Final pre-extraction internal standards concentrations are 50 μg/mL. Precellys 24 touch homogenization cycles were repeated and centrifuged as before. Homogenates were collected and pooled after both extractions. Metabolites were concentrated using a CentriVap Vacuum Concentrator at 30°C for 30-45 min and samples were stored at -80°C in the presence of argon gas until analysis by untargeted LC-MS. Metabolite peak signatures were exported as RAW (Thermo.RAW) files and imported into Compound Discoverer™ 3.5 software for metabolite identification. The metabolites were subsequently aligned and consolidated by positive and negative ionization modes. Peaks that corresponded to the same metabolite species were merged. The final list of metabolites and its abundances were imported into R for further analysis.

### Untargeted Liquid Chromatography and Mass Spectrometry

#### Proteins

Dried samples were reconstituted in 47 μL of 2% acetonitrile in water with 0.1% formic acid, 1 μL Trypsin-digested BSA MS Standard (New England BioLabs™, Ipswich, MA, USA; catalog no. P8108S), and 2 μL of Regen II human peptide standards A1315 (DRV[U-¹³C₅,¹⁵N]YI[U-¹³C₆,¹⁵N]HPFHL) and HH4B (SGRGKGGKGLGKGGAKRHRKVLRGGK-Biotin) were added at 54 μM each as post-extraction ionization controls. Both Regen III and Regen II peptide standards (>98% purity) were custom synthesized by AnaSpec (Fremont, CA, USA). After resuspension, the 18plex TMTpro labeled sample was sonicated in an ultrasonic water bath for 15 min for total solubilization. Samples were transferred to their respective autosampler vials. Untargeted liquid chromatography tandem mass spectrometry (LC-MS/MS) was performed on a ThermoScientific Vanquish Neo UHPLC System coupled to a QExactive Orbitrap mass spectrometer (ThermoFisher Scientific). The sample was ionized and detected using an EASY-Spray™ HPLC Column (catalog no. ES900, 150 mm x 75 µm x 3 µm, ThermoFisher Scientific) coupled to an EASY-Spray™ Source (catalog no. ES081, ThermoFisher Scientific) and 1 µL of sample was injected onto the column. The flow rate was 0.300 µL/min. Mobile phase A consisted of water with 0.1% formic acid (v/v) and mobile phase B was 80% acetonitrile in water with 0.1% formic acid (v/v). The column’s temperature was 55 ◦C. The spray voltage was set to 1.92 kV, and the capillary temperature was set to 300°C. The S-Lens RF level was set to 60.0. For full scans, the mass range was 375–1400 m/z, the resolution was 70,000, and there was 1 microscan. The AGC target was 3 × 106 and maximum injection time was 50 ms. For dd-MS2, the mass range was 200–2000 m/z, the resolution was 35,000, the AGC target was 1 × 10^5^, and the maximum injection time was 100 ms. The instrument was set to Top 10, the isolation window was set to 1.2 m/z, and the NCE (Normalized Collision Energy) was set to 32. The intensity threshold was 2.04 × 10^4^ and dynamic exclusion was 30s.

#### Lipids

Extracted lipids were dried and resuspended in LC-MS grade acetonitrile: isopropanol (1:1 v/v) and 1 μL of EquiSPLASH™ LIPIDOMIX® mixture (cat. no. 330731, Avanti Polar Lipids, Alabaster, AL). Following resuspension, samples were sonicated for 15 minutes for total solubilization. Reversed phase chromatography was performed on a Vanquish Horizon UHPLC system (ThermoFisher) coupled to an Accucore Vanquish C18+ UHPLC Column (Accucore™ Vanquish™ C18+ Column (150 mm × 2.1 mm, 1.5 μm, Thermo Scientific™, Waltham, MA, USA). An injection volume of 5 μL was used with a flow rate of 260 μL/min. Mobile phase A was 50% acetonitrile, 50% water, 5mM ammonium formate, and 0.1% formic acid. Mobile phase B was 88% isopropanol, 10% acetonitrile, 2% water, 5mM ammonium formate, and 0.1% formic acid. Ionization and detection were performed with a heated electrospray ionization source (HESI) coupled to a Q Exactive mass spectrometer. Data was collected in positive and negative mode for each sample. Spray voltage was 4.0kV in positive mode and 2.53kV in negative mode. For both modes: sheath gas flow rate was 35, auxiliary gas flow rate was 16, sweep gas was 0, capillary temperature was 325C, and S-lens RG level was 70. The full scan range was 250 to 1200 m/zm resolution was 70,000, and microscans was 1. AGC target was 1e6 and maximum injection time was 100ms. In dd-MS2, the resolution was 17,500, AGC target was 1e5, loop count was 10, and isolation window was 1.0m/x. NCE was set to 20,30,40, intensity threshold was 2.0e4, and dynamic exclusion was 8.0s.

#### Metabolites

Using untargeted LC-MS, samples were reconstituted in 44.75 µl LC-MS grade water and 0.1% formic acid, and post-extraction internal standards (all from Cayman chemicals): 0.25 μL of 5 mg/ml L-Phenylalanine-^13^C_6_ (catalog no. 35195), 2.5 μL of 0.5 mg/ml Uracil-^13^C,^15^N_2_ (catalog no. 33997), 1.25 μL of 1 mg/ml L-Arginine-13C6 (hydrochloride) (catalog no. 9004088), and 1.25 μL of 1 mg/ml L-Serine-^13^C_3_ (catalog no. 35126) to each sample. Concentrations of all post-extraction internal standards were 25 μg/mL. A small aliquot (10 μL) of each sample is pooled together in a separate reaction for quality control analysis by LC-MS to measure reproducibility and metabolite stability. Samples were run as technical replicates, undergoing fractionation and detection using a Thermo Scientific™ Vanquish™ Horizon Binary UHPLC system. Preparations of the mobile phase composition and separation by the QExactive™ mass spectrometer coupled to HESI source (Thermo Scientific™ Xcalibur™) have been detailed elsewhere.^22^ Briefly, an Accucore™ Vanquish™ HILIC Accucore Amide 150 Column (100 mm x 2.1 mm, 1.5 µm, Thermo Scientific) was used to separate compounds with a flow rate of 0.500 mL/min. For positive mode: Mobile Phase A consisted of 10 mM ammonium formate in 95% acetonitrile with 0.1% formic acid (v/v) and Mobile Phase B consisted of 10 mM ammonium formate in 50% acetonitrile with 0.1% formic acid (v/v). For negative mode: Mobile Phase A consisted of 10 mM ammonium acetate in 95% acetonitrile with 0.1% acetic acid (v/v) and Mobile Phase B consisted of 10 mM ammonium acetate in 50% acetonitrile with 0.1% acetic acid (v/v). Column temperature was set to 35 °C and injection volume was set at 5 μL. The spray voltage was set to 3.50 kV, capillary temperature to 350 °C, sheath gas to 55, aux gas to 14, sweep gas to 4, and S-Lens RF level to 30.0. The mass range was set to 67–1000 *m/z*; we set the resolution to 140,000 for full scans and to 35,000 for ddMS2. The AGC target was set to 1 × 10^6^ for full scans and 2 × 10^5^ for ddMS^2^. The max injection time (IT) was 100 seconds for full-scan mode and 50 seconds for ddMS^2^. The number of microscans was 2, and normalized collision energy (NCE) was set to 20, 35, and 50. Samples were run in both positive and negative ion modes, separately. The parameters for negative mode were the same with the exceptions of the spray voltage, which was set to 2.50 kV, and capillary temperature set to 380°C.

### Protein normalization and identification

The raw scans were processed via Proteome Discoverer 3.0.1<u>^TM^</u> (ThermoFisher Scientific) using a Sequest HT engine. The data were searched against *Homo Sapiens* entries (Swiss-Prot + TrEMBL, UniProt). The max missed cleavage site was set to 2 and the minimum peptide length was set to 6. Precursor mass tolerance was set to 10 ppm and fragment mass tolerance to 0.02 Da. Post-translational modifications for experimental proteins included oxidation, acetylation, carbamidomethylation, and TMTpro. Normalization was performed in relation to the total peptide amount. Target false discovery rate (FDR) was applied: 0.01 (strict) and 0.05 (relaxed). An 18plex TMT label reagent kit was used as a quantification method. Abundances were calculated after normalization with the TMTpro labels. A total of 13473 proteins were identified. To select for high-quality proteins, we filtered for candidates identified using at least 2 unique peptides or 1 unique peptide with an FDR rate of less than 5%. A total of 3141 protein candidates with high and medium confidence levels remained after the filter. Average abundances across technical replicates were calculated.

### Lipid identification

The alignment of positive and negative lipid species was individually performed with a Retention Tolerance of 0.05 min, Retention Correction Tolerance of 0.5 min, Signal to Noise Threshold of 3.0, Intensity Ration Threshold of 1.5, and Valid Peak Rate Threshold of 0.05. Lipid species were aligned by positive and negative mode and technical replicates consolidated. Lipid concentration was normalized to protein amount determined from the corresponding aqueous phase. The identified lipids were normalized by the deuterated internal standards. Lipids were standardized according to class-specific standards (1,2-ditridecanoyl-sn-glycero-3-phosphocholine, 1,2-dioleoyl-sn-glycero- 3-phospho-L-serine, phosphoserine, 1,2-dioleoyl-sn-glycero-3-phosphoethanolamine, and 1,2- dioleoyl-sn-glycero-3-phospho-(1’-myo-inositol).^23^

### Metabolite identification

The raw scans were processed via Compound Discoverer^TM^ 3.3. Positive and negative scans were analyzed for identification separately. Extraction blanks were used to determine and correct for reagent effects, mark background components, and filter background components from the results table. Pooled QCs were used for compound identification only. Metabolites with duplicate peak areas were merged and the compounds identified in both positive and negative mode were aligned.

### Data Analysis for proteomics, lipidomics, and metabolomics

Datasets were stratified into groups based on donor characteristics (age and sex) and media conditions (high vs low glucose media and laboratory source) and analyzed using R statistical analytical packages (pheatmap, ggplot2, tidyverse, UpSetR).^24–27^ For heatmap analysis, log transformation (base 2) of the abundances was performed across all samples and z scores calculated. Hierarchical clustering by Euclidean distance and Ward’s linkage method were applied to the dendrograms according to age, sex, laboratory, and media conditions. Principle component analysis and volcano plots were generated by tidyverse R packages for comparing group characteristics. When specified, P-adjusted values utilized the Benjamini-Hochberg correction with FDR <0.05.

#### Technical validation

The datasets generated herein will be useful to glaucoma researchers and clinicians, who are interested in comparing their data with ‘normal’ TM cells. This includes metanalyses, preclinical drug discovery studies, researchers interested in comparing data from glaucoma animal models, etc. It is also useful for researchers investigating specific parameters. For example, we have provided comparisons of how age, sex, and glucose impact each of the omics datasets. Transcriptomic data can also be compared to single-cell RNAseq datasets.^28–30^

##### Characterization of TM Cell Strains

All TM cells were characterized by standard dexamethasone induction of myocilin^31^ by either Western blot (**Figure 2A**), or quantitative RT-PCR analysis (**Figure 2B**).

**Figure 2.**
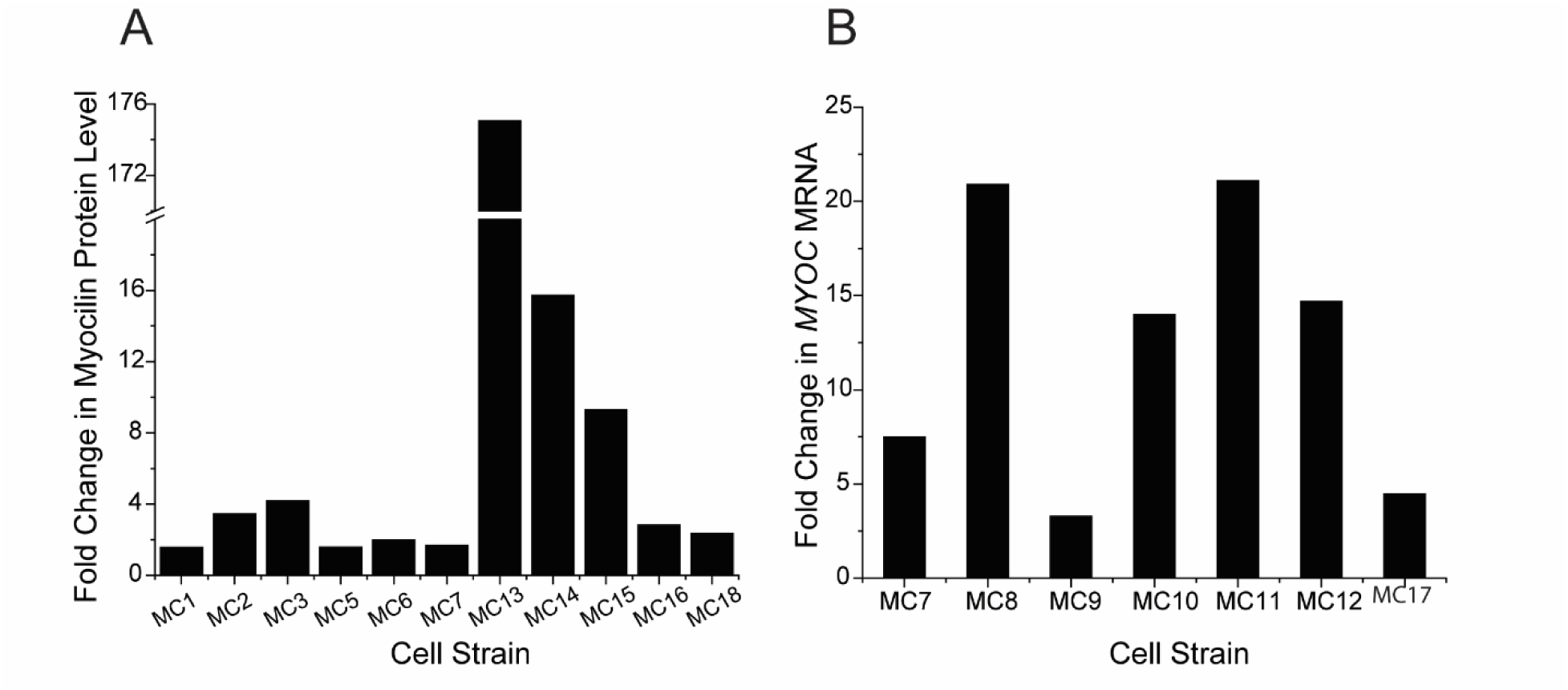
Validation of TM cells by upregulation of myocilin following treatment with 100nM dexamethasone for 5-8 days. (A) Fold increase in myocilin protein levels over ethanol control detected by Western blot analysis, normalized to GAPDH (1-3, 5-7, 15, 16, and 18) or ponceau S (13, 14). (B) Fold increases in mRNA levels were normalized to the SDHA or GAPDH housekeeping genes (4, 8, 9, 10, 11, 12, 17).

For metabolomics, three technical replicates of each sample were normalized against six types of standards (arginine, glucose, isoleucine, phenylalanine, serine, and uracil). The coefficient of variation (CV) percentage was calculated across technical replicates for each sample (**Figure 3A**), and across each normative standard is shown in a graph (**Figure 3B**). Most samples had a CV of less than 30% CV across multiple normative standards. The MC3 sample exhibited the highest CV% in the phenylalanine standardization. Arginine, glucose and uracil standardization yielded a wider range of CVs (24-66%, 19-57%, and 20-70%, respectively) than isoleucine and serine (20-35% and 23-54%, respectively). Over 70% of detected metabolites showed a CV of less than 40% in five of seven laboratories (**Figure 3B**). Laboratory 4 showed a higher CV percentage only in the arginine, glucose, phenylalanine, and uracil standardizations. Laboratory 7 showed a higher CV percentage during the standardization of arginine, glucose, and uracil.

**Figure 3.**
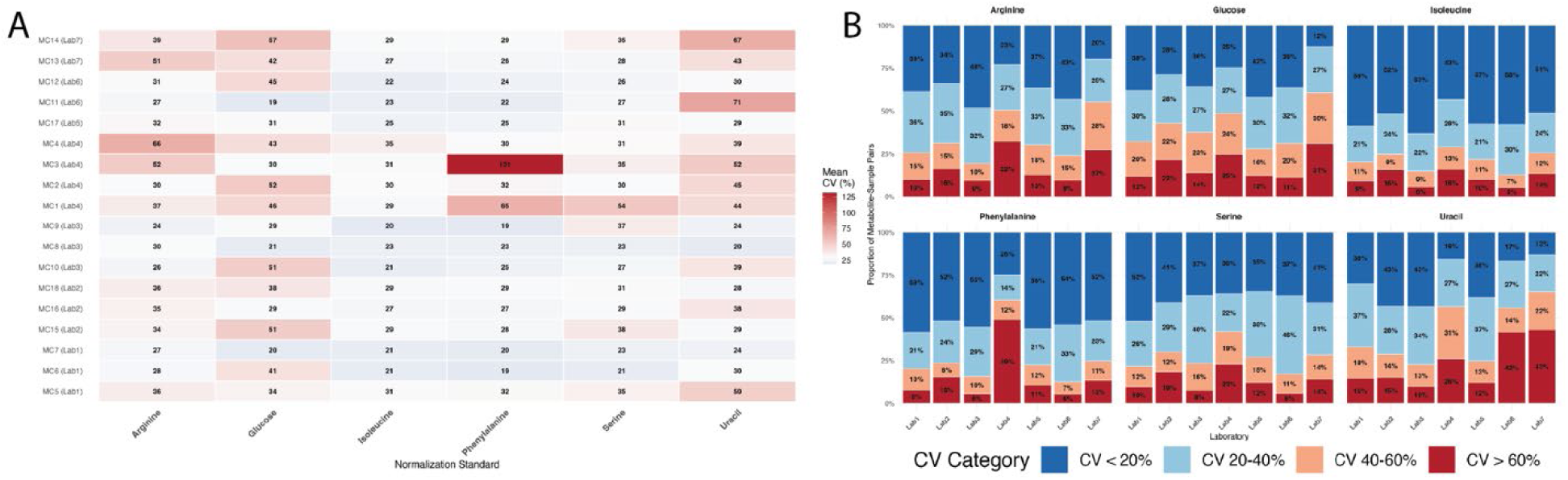
Coefficient of variation (CV) analysis of metabolome. Matrix of CV (%) averaged across technical triplicates for 18 TM cell strains (A). Proportion of detected metabolites with stratified CV (%) that is graphed by each contributing laboratory and across each normative standard (B).

For all ‘omics, group comparisons of the data by age, sex, and glucose concentration in the media were performed. For age, the 18 cell strains were separated into 3 age groups: < 40 years (n=4), 41- 59 years (n=7), and > 60 years (n=7) (**Table 1**). For sex, there were n=7 females and n=11 males (**Table 1**). For glucose, n=10 cells were cultured in low-glucose (1000 mg/L) media, while n=8 were cultured in high-glucose (4500 mg/L) media (**Table 2**).

### Transcriptomics analysis

*RNA integrity assessment.* Assessment of RNA quality and concentration was conducted with a NanoDrop 2000 spectrophotometer and Agilent 2100 Bioanalyzer. RINs were relatively low (<7), but the values have been shown to be reliable for RNAseq.^32^ After quality control of RNAseq raw data, an average of 98.99 million high-quality reads per sample (range: 26.9-207.8M) were obtained (**Table 3**). The alignment statistics to the reference genome are also summarized in the table.

**Table 3.** RNA integrity assessment for the 17 TM cell strains.

| Sample ID | Read Pairs | Transcriptome Aligned |
| --- | --- | --- |
| MC1 | 243,455,459 | 130,417,914 |
| MC2 | 207,772,764 | 73,690,532 |
| MC3 | 115,584,663 | 65,227,566 |
| MC5 | 84,922,735 | 53,653,824 |
| MC6 | 37,967,262 | 27,659,970 |
| MC7 | 70,144,248 | 43,992,996 |
| MC8 | 55,623,202 | 29,602,159 |
| MC9 | 160,564,891 | 81,516,464 |
| MC10 | 63,288,901 | 33,474,881 |
| MC11 | 55,424,269 | 23,669,690 |
| MC12 | 137,238,722 | 66,243,469 |
| MC13 | 50,686,940 | 30,630,166 |
| MC14 | 105,430,114 | 56,979,650 |
| MC15 | 24,906,257 | 16,158,632 |
| MC16 | 161,954,103 | 89,676,365 |
| MC17 | 81,329,612 | 50,736,332 |
| MC18 | 64,336,462 | 64,336,462 |

A Principal Component Analysis (PCA) plot was used to visualize the similarity of transcriptional profiles of cell strains derived from independent laboratories (**Figure 4A**). This analysis indicates that biological variation between cell samples exceeds the technical variation due to minor differences in culture conditions. Additionally, a heatmap including the top 100 most variable genes was used to identify gene expression patterns correlated with potential experimental factors of interest (laboratory, age, sex, media glucose level) (**Figure 4B**). Despite high variability across cell strains, this variation was not strongly associated with any of these experimental factors. However, a small number of genes were significantly upregulated or downregulated in each group comparison are shown in **Figure 4C**.

**Figure 4.**
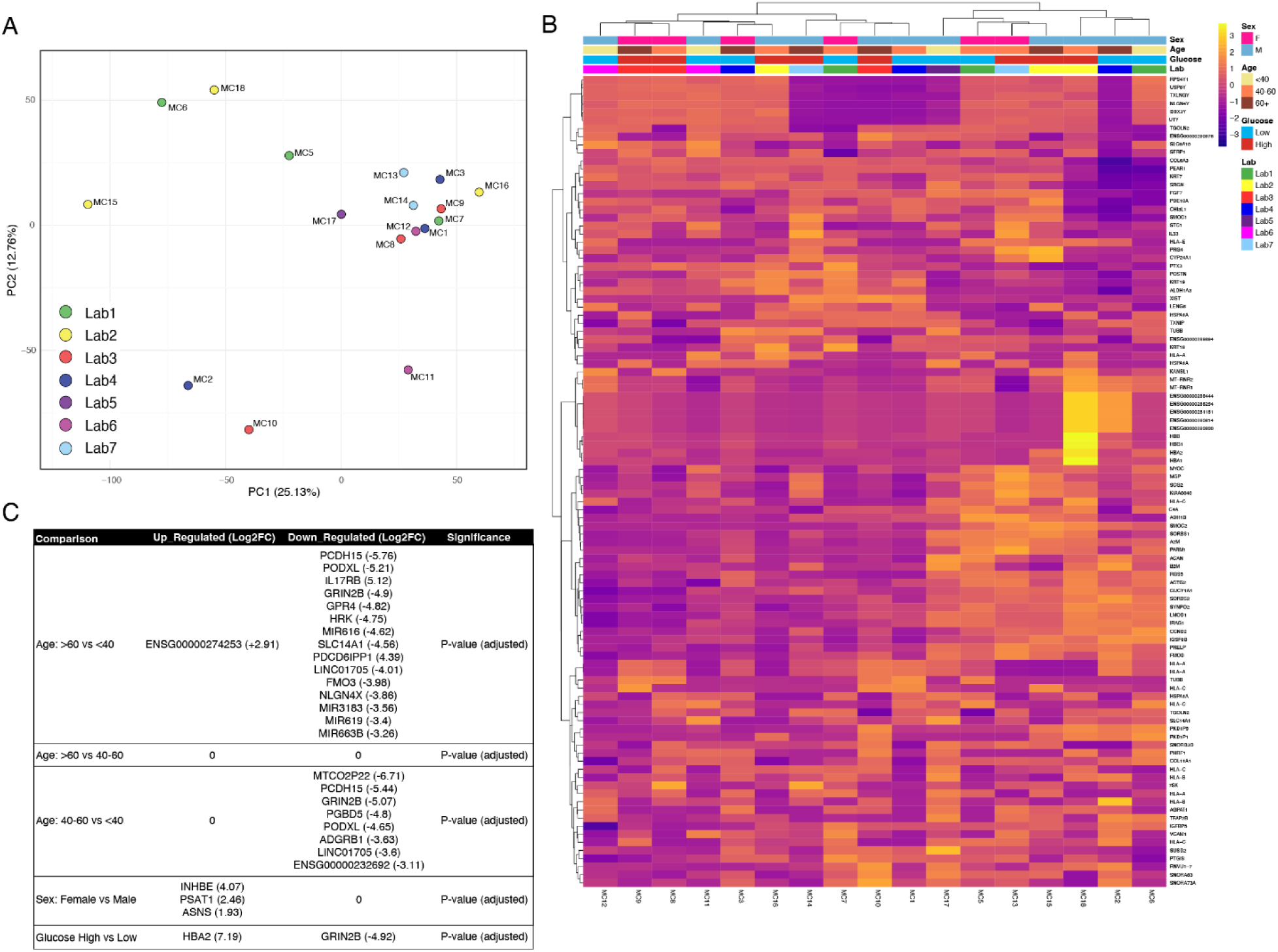
Bulk transcriptome sequencing. (A) PCA analysis was done to show the similarity between TM cell strains. The closer the cell strains, the higher their similarity. The color indicates the laboratory that isolated the TM cell strain. (B) Heat maps of all the transcripts were generated according to laboratory, age, sex, level of glucose in tissue culture medium used. (C) A table of genes with significant differential expression between comparative groups.

### Proteomics analysis

Each of the 18 TM cell strains were labeled with a unique tag for peptide identification and normalized abundances. Among the total of 13473 proteins identified, application of quality filters provided 3141 high- and medium-confidence candidates for analysis (see Methods). The number of unique proteins identified are comparable to past proteomic studies of human TM cell cultures.^33,34^ The PCA plot demonstrated one large cluster containing a majority of strains from all the depositing laboratories (**Figure 5A**). Two strains, MC9 and MC6, showed highest variability along PC1 and PC2, respectively. The diversity of these strains likely reflects intrinsic proteomic differences in the TM donor rather than experimental factors (laboratory source or glucose condition) since these strains clustered differently in the lipidomic and transcriptomic analyses. Heatmap analysis showed moderate variability within the major cluster (**Figure 5B**). Similar to the transcriptomic analysis, there were only a few proteins with small expression changes after grouping by biological factors (age and sex) and media condition (high vs low glucose) (**Figure 5C**).

**Figure 5.**
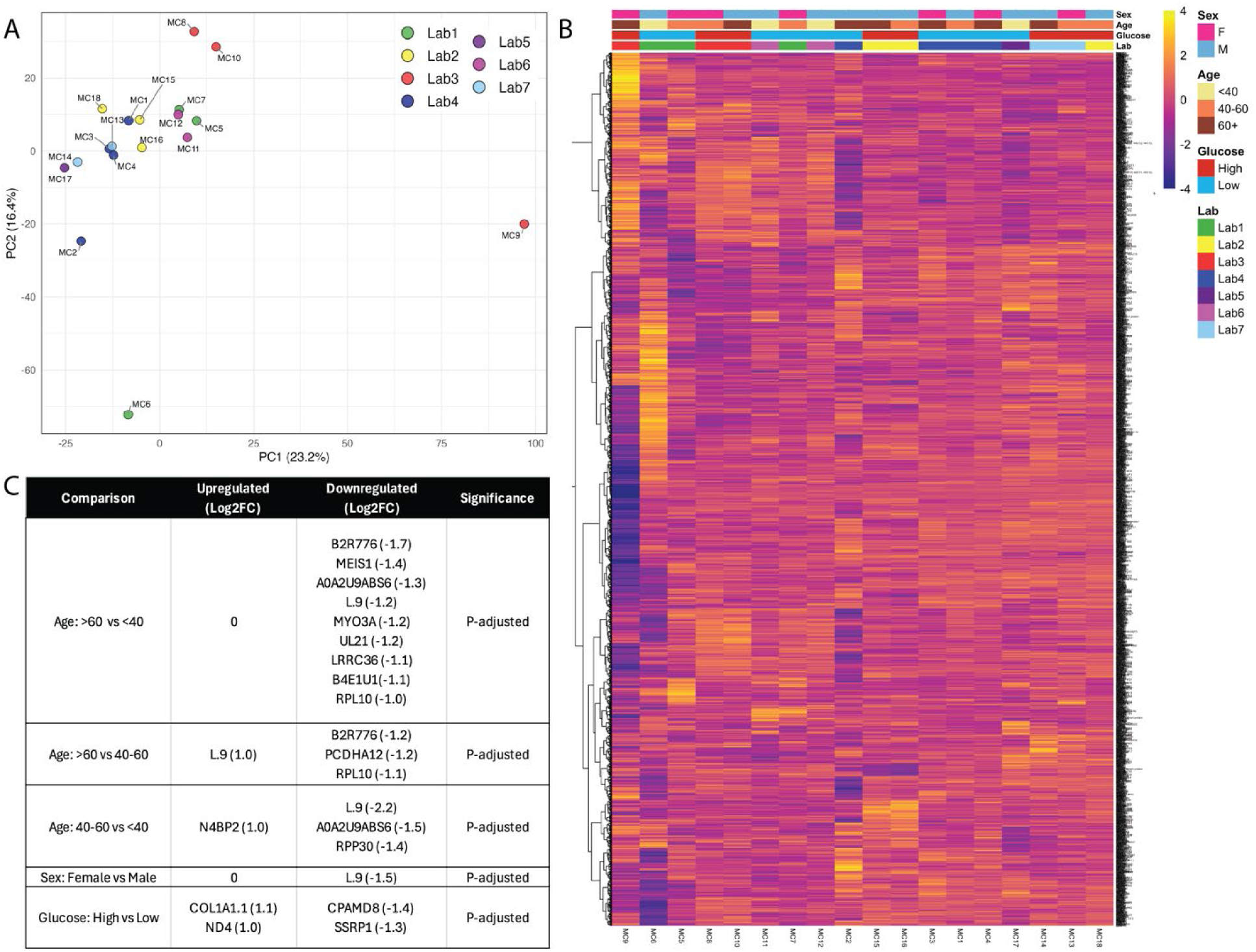
Summary of proteomic dataset including PCA analysis (A), heatmap (B), and a table of proteins (C) differentially expressed between comparative groups. P-adjusted values utilized the Benjamini-Hochberg correction with FDR <0.05.

### Lipidomics analysis

There were two distinct clusters in the lipidome dataset based on depositing laboratory (**Figure 6A**). These clusters were maintained for all of the comparisons (age, sex, glucose concentration in media, **Figure 6B**) and for pairwise laboratory comparisons using Jaccard similarity indices (**Figure 6C**). Heatmap data and Jaccard similarity indices also clearly showed the clustering into two groups (**Figure 6B and 6C**).

**Figure 6.**
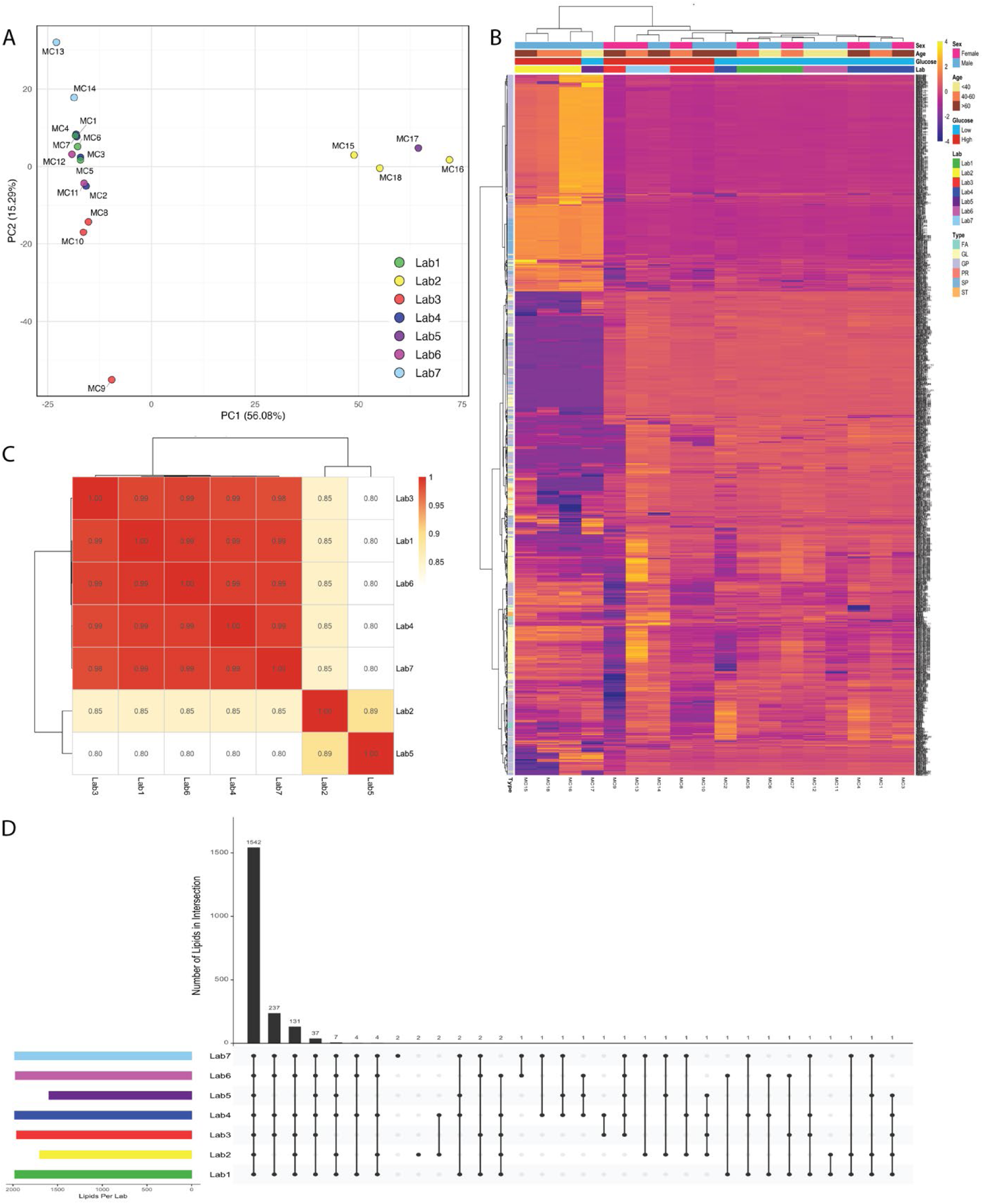
Summary of lipidomic data. PCA analysis (A) and heatmap (B) for all samples and characteristics. Analysis of total lipid types detected from individual laboratories shown as heatmap of Jaccard similarity index (C). UpSet matrix graph (D) showing common lipid types identified across different laboratories.

Cluster 1 comprised the majority of cell strains with a total of 14 TM cell strains (MC1-14). The TM cells came from five separate laboratories and were derived from donor eyes from individuals aged 25-74 years. The lipid profile of this cluster consisted of four triglycerides (TG) and 6 phosphatidylethanolamines (PE). Together, they constituted the majority of the lipids found in these cells. Two diglycerides (DG), two phosphatidylglycerols (PG), and a hexosylceramide (Hexcer) were also detected. This cluster is considered the main lipidomic profile of TM cells, as it contains the majority of TM cells.

Cluster 2 is smaller and contains 4 cell strains (MC15-18). Three of these came from one laboratory, which used media supplemented with GlutaMAX. Research has shown that GlutaMAX can reduce triglycerides and other lipid levels. This smaller cluster consisted of cell strains derived from individuals aged 29-83. The lipid profile of this cluster of TM cell strains differed from the larger cluster with decreased PE lipids, but increased sphingomyelins and ceramides.

Although there appears to be two distinct clusters in the lipidomic analysis, it is important to note that there are remarkable similarities across all samples. Analysis of total lipid types detected showed that 77% of lipid species were observed in all laboratories. Another 9% and 12.0% of unique lipid species were identified in 6 of 7 and 5 of 7 laboratories, respectively (**Figure 6D**).

To further assess the quality of the data, we examined the effects of glucose levels on the lipid profiles of TM cells (**Figure 7A**). TM cells cultured in high glucose had upregulation of 8 glycerolipids (GL) and 1 fatty acyl (FA) lipid, while there were 50 lipids that were more enriched in the low glucose group (Log2FC ≥ ±1; p<0.05). These were predominantly GPs and SPs. Glucose did not affect prenol or sterol lipid profiles. Sex also influenced lipid profiles (**Figure 7B**). GL (n=13) were upregulated in females, while glycerophospholipids (GP) (n=15) and sphingolipids (SP) (n=35) were more enriched in males (Log2FC ≥ ±1; p<0.05). There was n=1893 lipids that showed no sex differences. Sex did not significantly affect fatty acids, prenol, or sterol lipid profiles.

**Figure 7.**
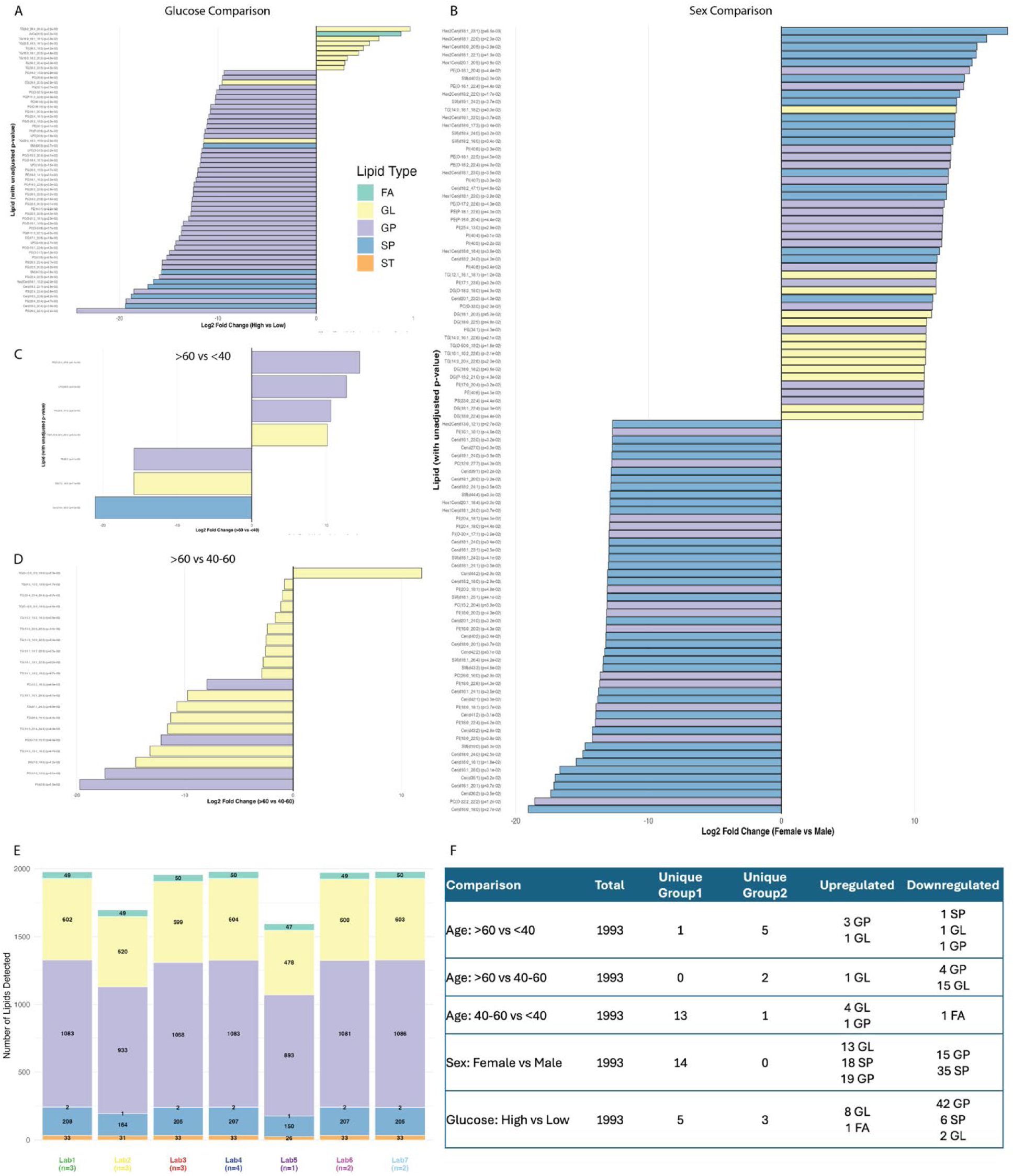
Lipidomic analysis by glucose levels in media (A), sex (B), age comparisons (C, D), and laboratories (E). Summary table (F) of total unique lipids and significantly upregulated and downregulated lipid type across all comparative analyses. Fatty acids (FA); glycerolipids (GL), and glycophospholipids (GP).

There was a significant age effect on lipid profile (**Figure 7C**). This included significant decreases in triglycerides in TM cells derived from older individuals compared to the middle-aged (41-59 yrs) group (Log2FC ≥ ±1; p<0.05). This comparison yielded the largest number of lipids (n=20) showing differential levels and may be important because individuals >60 years^35^ are more likely to develop glaucoma than the middle-aged group. The lipid profile of cells from individuals aged 41-60 also showed some differences compared with that of younger donor TM cells. These included an increase in 4 TGs and 1 DG. Most lipids (n=1973) did not show age-related changes to their expression profiles. Age did not significantly affect prenol or sterol lipid profiles.

### Metabolomics analysis

A PCA plot and heatmap were generated from metabolomics data normalized by arginine (**Figure 8A, B**). These show two clusters each composed of multiple depositing laboratories: cluster 1: MC1- MC5, MC13-MC18; and cluster 2: MC6-MC12. Data was also analyzed by normalization to isoleucine (**Figure 8C**), which showed similar clusters, although the PC2 gave different values for some of the samples in cluster 1. Similarly, data was normalized to glucose, uracil, phenylalanine, and serine. A table showing the total number of metabolites (n=254) defining the PC1 and PC2 variance for each of the normative standard is shown (**Figure 8D**).

**Figure 8.**
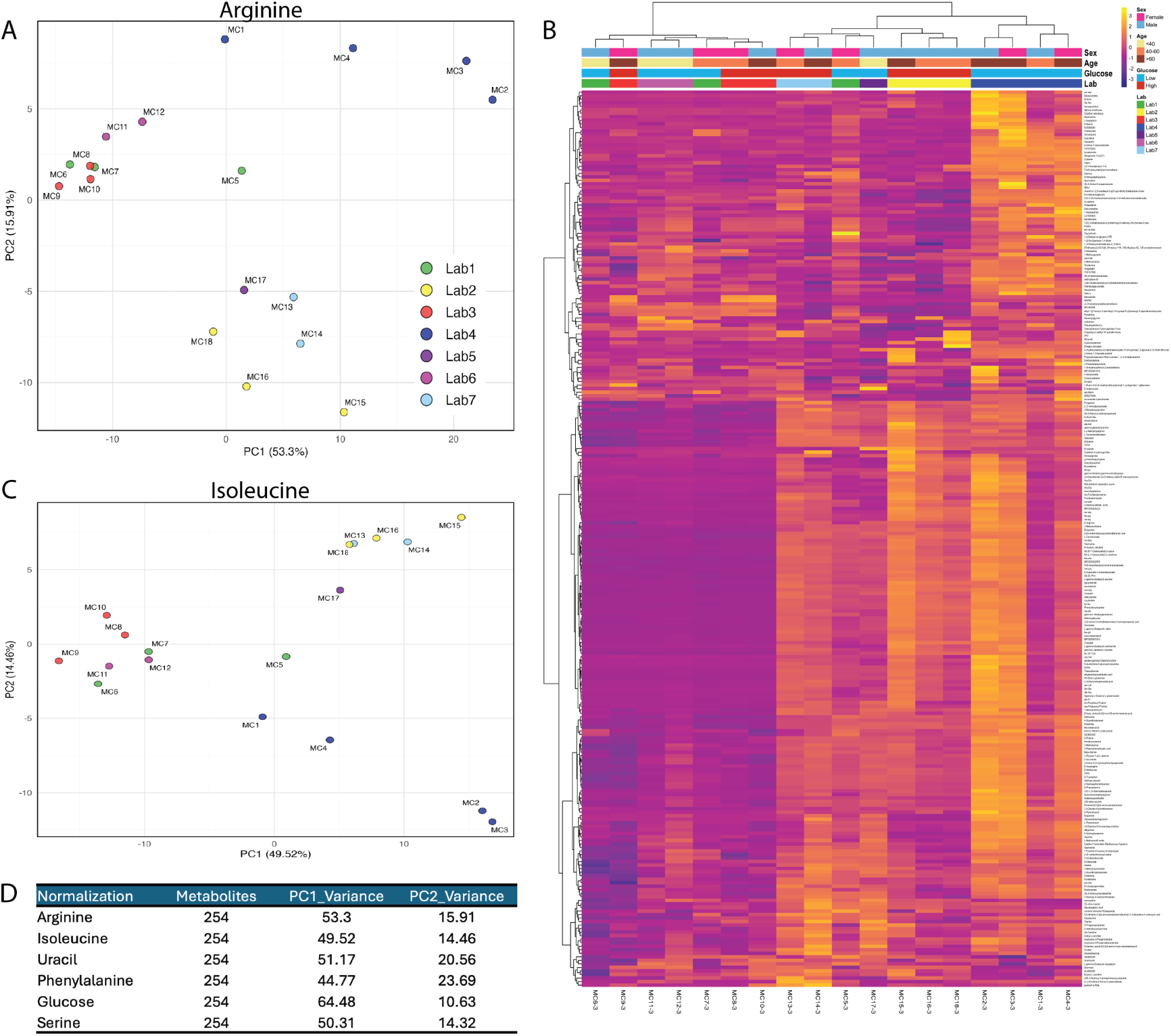
Metabolomic PCA and heat map analysis. PCA analysis from normalization with arginine (A, B) or isoleucine (C) standards are clustered by depositing laboratories. (D) Table summarizes the variance across all normative standards.

To assess the quality of the data, we examined the top 7 upregulated metabolites and found that they were the same regardless of normalization method (spermine, HEPES, gauaiacol sulfate, butyryl-L-carnitine, AL9450000, Minoxidil and (2R)-2-hydroxy-3-(phosphonooxy)-propanal). Similarly, the top 5 most downregulated are the same regardless of normalization method (Cyclidine- 1-beta-ribofuronosyl-cytosine, guanine, ubiquinon-1(CoQ1), TX1575000, Inspra®). Many of the metabolites detected were drugs or chemicals. Systemic drugs such as blood pressure medication (Inspra®), anti-anxiety (Buspirone, Imagabalin), hair loss (Minoxidil), treatment for gout and to prevent kidney stones (Allopurinol), and anti-seizure medication (Etiracetam) were found. Environmental chemicals detected included cetonecyanohydin (a key component of plexiglass) and HEPES, which was high in the samples that were cultured in DMEM/F12 (samples MC8-10). These various drugs and chemicals were detected in TM cells even after multiple passages.

Metabolites were analyzed by age, glucose concentration, or sex (**Figure 9**). A dot plot shows comparisons between the >60 and the <40 age groups for each normalization. The color of each circle delineates up- (red) or downregulated (blue), while the size of the circle determines the significance (log10 p-value) of metabolite enrichment. Most metabolites were upregulated in TM cells derived from older individuals (**Figure 9A**). Normalization by uracil yielded a pattern that was most dissimilar to the other normalization patterns. The increase in many metabolites (17 of the top 50) observed in the older group is attributable to exogenous metabolites derived from medications or environmental factors, suggesting these affected some cells, but not others. Data were analyzed by the glucose concentration (high vs low) (**Figure 9B**). When normalized to arginine, there were more downregulated metabolites than upregulated. The opposite was true when normalizing to uracil. Sex did not impact metabolites as much as age or glucose as there were fewer metabolites in this comparison (**Figure 9C**). Normalization by glucose and phenylalanine gave the most dissimilar patterns in this sex comparison. The total number of metabolites, as well as how many were up- or downregulated, for each comparison (age, sex, glucose concentration) is also shown (**Figure 9D**). The number of up- (**Figure 9E**) or downregulated (**Figure 9F**) metabolites detected in each of the comparisons by normalization procedure are also shown.

**Figure 9.**
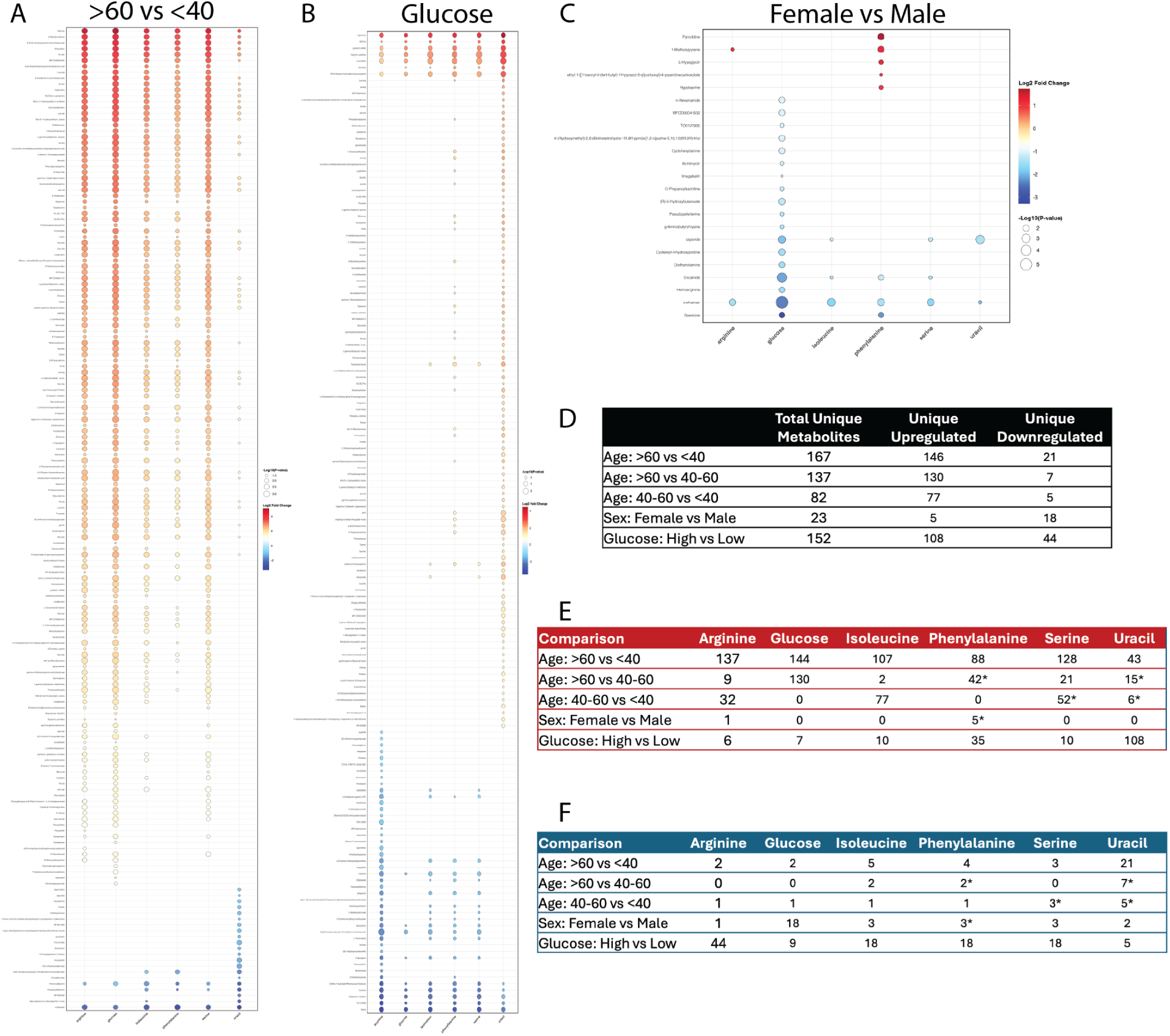
Metabolomics dot plot for age (A), media (B), and sex (C) comparison. Tables D-F lists number of unique metabolites in total (D), upregulated (E), or downregulated (F) across different comparative analyses. Red represents upregulated and blue represents downregulated metabolites. Size of dots correlate with size of log10(p-value). *P-unadjusted used for comparative analysis. When not specified, p-adjusted value used elsewhere. P-adjusted utilized Benjamini-Hochberg correction with FDR < 0.05.

## Code availability

All R codes used are detailed in the methods and are available upon request.

## Acknowledgements

The authors appreciate the support of Miami metabolomics research support group (https://www.mmrsg.org/). The authors would also like to thank the eye banks that procured and made available the eye tissue for use in this study: VisionGift, Portland, OR; Lions Eye Bank of Wisconsin, Madison, WI, Liverpool Research Eye Bank, UK; Beauty of Sight Eye Bank, FL; and LIONS World Vision Institute, FL.

This research was supported by NIH grants R01-EY031292 (SKB), R01-EY032590 (KEK), R01- EY019643 (KEK), R01-EY17006 (DMP), R01-EY032905 (DMP), R01-EY034096 (SH), R21-EY036189 (SH), R01-EY025643 (YD), R01-EY036983 (YD), R01-EY035412 (PPP), R01-EY029320 (PPP), K08- EY037401 (AHP), P30-EY016665 (DMP), P30-EY010572 (KEK), and P30-EY014801 (SKB, AHP). The research in the Swaroop laboratory is supported by the Intramural Research program of the National Eye Institute (ZIAEY000450, ZIAEY000546) and utilized the computational resources of the NIH HPC Biowulf cluster (https://hpc.nih.gov). The contributions of the NIH authors are considered Works of the United States Government. The findings and conclusions presented in this paper are those of the authors and do not necessarily reflect the views of the NIH or the U.S. Department of Health and Human Services. The research presented was also supported in part by: Unrestricted grants from Research to Prevent Blindness (New York, NY) to the Casey Eye Institute, OHSU (KEK) and to the Department of Ophthalmology & Visual Sciences, University of Wisconsin-SMPH (DMP), the Department of Ophthalmology and Visual Sciences, Center for Vision Research at SUNY Upstate Medical University (SH), the Glick Eye Institute, Indiana University School of Medicine (PPP), and Bascom Palmer Eye Institute (AHP, SKB); Glaucoma UK (CMS); American Glaucoma Society Early Physician Scientist Award (AHP), Stanley J. Glaser Foundation Award (AHP); BrightFocus Foundation National Glaucoma Research Award G2024006S (SH); Ralph W. and Grace M. Showalter Research Trust (PPP), and Research Support Funds Grant (RSFG) Indiana University School of Medicine (PPP).

## Contributions

Conceptualized and designed the study: SKB, PPP, CMS, YD, NV, DMP, AHP, KEK.

Funding acquisition and Supervision: SKB, KEK, DMP, AHP, PP, CS, YD, SH, ASw.

Experimental Investigation: IGM, SKB, RPK, SL, EB, NS, MAE, JAF, YYS, ASo, RD, SP.

Formal analysis: ZB, NS, AHP, KK, IGM, TN, SHo, ASo.

Writing – original draft: AHP, KEK, DMP.

Writing – review and editing: All authors reviewed and edited the manuscript.

